# Live cell imaging of meiosis in *Arabidopsis thaliana* - a landmark system

**DOI:** 10.1101/446922

**Authors:** Maria Ada Prusicki, Emma Mathilde Keizer, Rik Peter van Rosmalen, Shinichiro Komaki, Felix Seifert, Katja Müller, Erik Wijnker, Christian Fleck, Arp Schnittger

## Abstract

Meiosis is essential for sexual reproduction and key to the generation of genetic diversity. To reveal the robustness of meiocyte differentiation and progression through meiosis, we have here established a live cell imaging setup to follow the dynamics of individual male meiocytes in Arabidopsis. Our method is based on the concomitant visualization of microtubules and a meiotic cohesion subunit that allowed following five cellular parameters: cell shape, nucleus position, nucleolus position, chromatin condensation and microtubule array. We find that the states of these parameters are not randomly associated and identify 11 states, referred to as landmarks, that occur much more frequently than closely related states, indicating that they are convergent points of meiotic progression. With this, the here-presented landmark system represents a novel method to analyze meiosis not only allowing a high-temporal dissection but also providing new criteria to evaluate mutants or environmental effects on meiosis.

## Introduction

Meiosis is essential for sexual reproduction by reducing chromosome number to eventually generate gametes with half the genomic DNA content as the parental plant. Moreover, meiosis is central to the formation of genetic diversity by generating recombination between the homologous chromosomes (homologs) and by randomly selecting either the maternal or paternal homologs into a new set of chromosomes in the gametes. Hence, understanding the molecular mechanisms underlying recombination as well as chromosome distribution and subsequent modulation of meiosis are also of key interest for breeding (Crismani et al., 2013; Hand and Koltunow, 2014; Lambing and Heckmann, 2018).

Entry into meiosis is tightly regulated in all organisms. In plants, it involves the reprogramming of somatic fate since plants, in contrast to animals, do not have a germline that is set aside early during embryo development (Schmidt et al., 2015). Designated meiocytes adopt a characteristic shape that radically changes during the course of meiosis ultimately resulting in the formation of spores. These spores then differentiate into gametophytes that produce the gametes, which will fuse during fertilization.

In recent years, our understanding of meiosis in plants has been fostered by genetic approaches, mostly in the model plants *Arabidopsis thaliana*, *Zea mays* and *Oryza sativa*. These studies have now identified more than 80 meiotic genes, including those that control entry and progression through meiosis (Lambing et al., 2017; Ma, 2006; Mercier et al., 2015; Wijnker and Schnittger, 2013; Zhou and Pawlowski, 2014). However, cytological studies of mutants defective in these genes have so far exclusively relied on the analysis of fixed material by cytochemical methods such as chromosome spreads and the immuno-detection of proteins. While these techniques have been, and continue to be, very informative, they capture the underlying cellular dynamics only to a small degree. Importantly, these methods do not allow individual cells to be followed over time. Thus, conclusions about the course of meiocyte development and progression through meiosis have to be deduced from the analysis of different cell populations at different time points.

So far, only two approaches to observe meiosis in real time in plants have been described, revealing details about spindle dynamics and chromosome paring in maize meiocytes. First, the work of Yu et al. and its modification by Nannas et al. used fluorescence microscopy to observe isolated male meiocytes cultured for a maximum of 9 hours in liquid medium (Nannas et al., 2016; Yu et al., 1997). The method of Nannas et al. combined the DNA dye Syto12 with the expression of β-tubulin fused to CFP, thereby allowing the concomitant observation of chromosomes and microtubules. This revealed spatially asymmetric distribution of the spindle at anaphase I and II, and chromosome-depending phragmoplast deposition (Nannas et al., 2016). The second approach involved imaging entire anthers of maize by exploiting the high depth of field of two-photon microscopy, as earlier proposed by Feijó et al. (Sheehan and Pawlowski, 2012, 2009). This method, which allowed imaging for periods of 24 hours, led to the characterization of three different movements and trajectories followed by the chromosomes during pairing in prophase I (Sheehan and Pawlowski, 2009).

These studies in maize relied on visualizing DNA by chemical stains such as Syto12 and DAPI and the power of Arabidopsis as a molecular model, which enables the relatively fast generation of fluorescent reporter lines for different meiotic proteins, has largely not been exploited in combination with live cell imaging of meiosis. A first approach was made by Ingouff et al. who observed methylation changes during Arabidopsis sporogenesis and gametogenesis, albeit without resolving specific meiotic stages (Ingouff et al., 2017).

Here we set out to develop a live cell imaging system for meiosis in Arabidopsis. To this end, we have generated an easy applicable microscopic set up, a combination of meiotic reporter lines covering central aspects of meiosis, and an evaluation system based on morphological characteristics that allowed the quantification of meiotic phases with high temporal resolution. This work gives insights into the robustness of meiocyte differentiation steps and provides important criteria to judge and/or re-evaluate mutants affecting meiosis.

## Results

### Specimen preparation

Live cell imaging can be performed at three general levels and all three have been applied to the analysis of meiosis in multicellular organisms. First, imaging can be performed on isolated cells as for instance seen in the case of mammalian oocytes (Holubcová et al., 2013; Kitajima et al., 2011; Kyogoku and Kitajima, 2017; Schuh and Ellenberg, 2007). This approach usually gives very high spatio-temporal resolution since there is no requirement for the laser beam to penetrate surrounding cells. Additionally, very little laser power can be used reducing photobleaching and phototoxicity. However, since meiocytes in Arabidopsis are very small i.e. 20 μm, and difficult to isolate, we did not explore this possibility further. Next, imaging can be carried out in the context of an entire organism, e.g. in *C. elegans* (Mullen and Wignall, 2017; Rosu and Cohen-Fix, 2017) with the benefit of perturbing the analyzed cells as little as possible by preparation procedures. However, this set up is limited to small organisms and/or short observation times due to size restrictions and the problem of movement of the sample, e.g. the elongated stem that carries the flowers in Arabidopsis results in a high degree of instability of the specimen during image acquisition. Hence, such a set up was also excluded. Finally, live cell imaging can be performed on isolated organs or tissues that are typically easy to obtain and that provide the appropriate developmental context for analysis of the selected individual cells, e.g. in mice (Enguita-Marruedo et al., 2018), *C. elegans* (Mlynarczyk-Evans and Villeneuve, 2017) and *Drosophila melanogaster* (Głuszek et al., 2015). As conventional confocal laser scanning microscopes can reach cells up to a depth of 70-100 μm, they are suited to observe the meiocytes in Arabidopsis that are covered by three cell layers in the anthers. Imaging of isolated organs has already been successfully applied to the analysis of organogenesis in the shoot apical meristem (SAM) of Arabidopsis (Hamant et al., 2014). Since shoots could be maintained for several days without obvious perturbations of development, we decided to adapt and optimize this approach for our purposes.

First, we selected inflorescences and removed all but one young flower primordium presumably containing meiotic stages as indicated by its round shape and an approximate diameter of 0.4-0.6 mm (Figure 1B), corresponding to stage 9 of flower development (Smyth et al., 1990). Next, the upper sepal was removed giving access to two of the six anthers since the petals are shorter than the anthers at this floral stage. Finally, the bud along with the pedicel and a few millimeters of the stem was embedded into Arabidopsis Apex Culture Medium (ACM) and stabilized with a drop of agarose (Figure 1A,B). In agreement with the previous analysis of the SAM, we found that the flower buds stayed alive on the ACM medium for up to seven days during which flowers grew and developed normally (Figure 1C).

**Figure 1.**
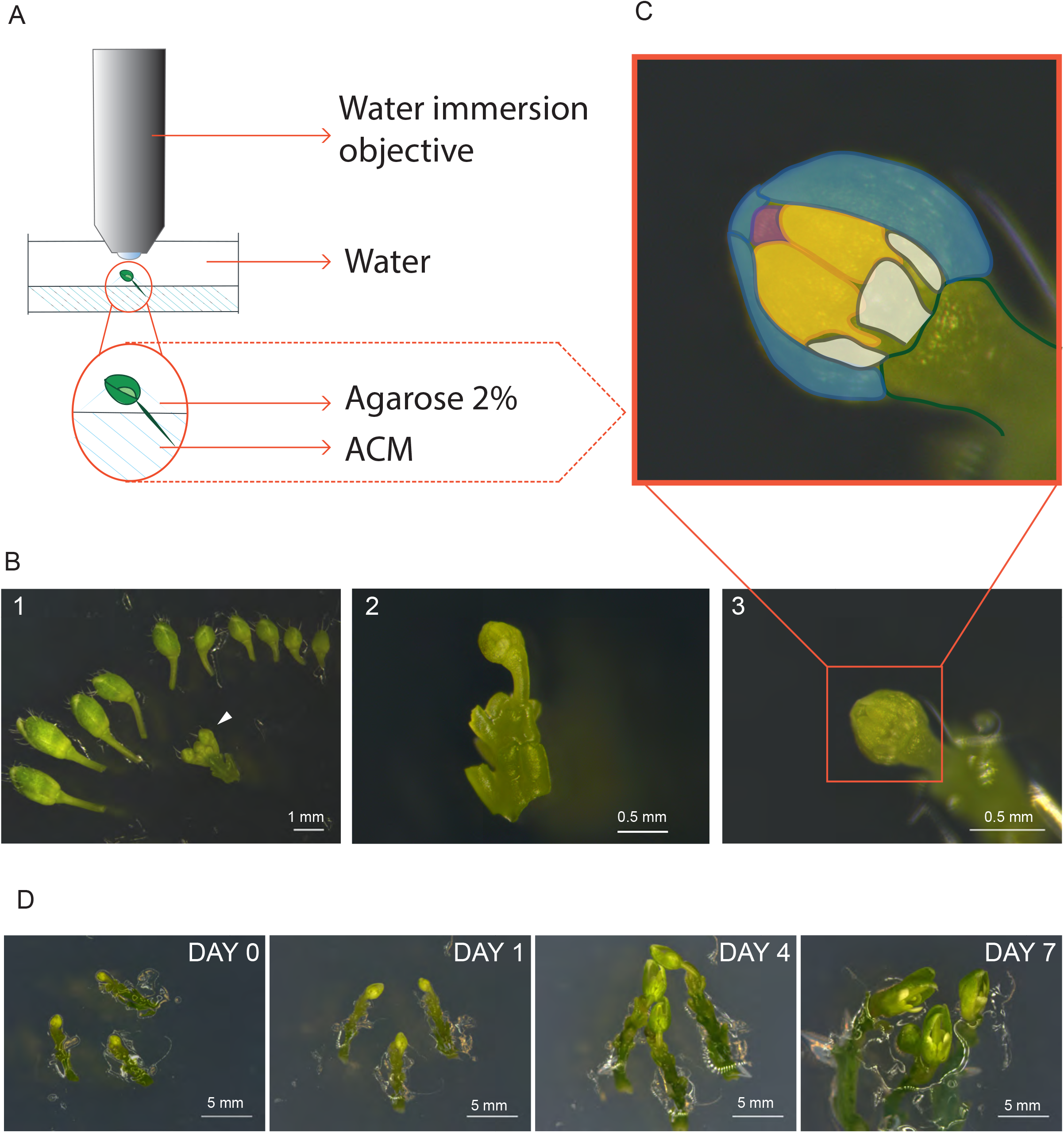
Technique establishment. A) Microscope set-up for live cell imaging. An isolated flower bud is mounted on a small petri dish in ACM medium, stabilized with a drop of 2% agarose and submerged with sterile water. The objective is directly dipped into the water. B) Steps of sample preparation. A flower bud of 0.4-0.6 mm length is selected (white arrow-head) (B1). The upper-most sepal of this flower as well as all other flowers are removed (B2). The flower is anchored into the medium with the anthers exposed at the top (B3). C) Magnification of the sample from B3. The two exposed anthers are highlighted in yellow; petals are in white, the three remaining sepals in blue, and the tip of the stigma in pink. D) Flower buds could be kept alive and growing for up to 1 week.

Imaging was performed with an up-right confocal laser scanning microscopy equipped with a water immersion objective. The entire flower bud was submerged in water and the objective was brought into direct contact with the sample (Figure 1A). During image acquisition the temperature was kept constant at 21°C.

### Establishment of meiotic reporter lines

A generic set up for imaging of cell divisions includes a reporter that highlights DNA/chromatin coupled with a marker for cytoskeletal components, usually microtubules, so that chromosome and spindle behavior can be visualized (Nannas et al., 2016; Peirson et al., 1997). Since fusions of histones with fluorescent proteins have often been applied for this purpose, we first scanned through previously generated transgenic lines expressing different histone variants fused to fluorescent proteins, such H2B. However, while these labeled histones clearly marked DNA in somatic cells, the signal was often fuzzy in meiosis. Moreover, since all or most cells in an anther produced these fusion proteins, the identification of meiocytes was sometimes difficult, especially at early stages of meiosis when chromosomes are not condensed and meiocytes cannot easily be recognized by their size and shape. Therefore, we aimed for a meiosis-specific gene and generated a *GFP* fusion to *REC8*, the alpha kleisin subunit of the cohesin complex, also known as *SYN1* or *DIF1* in Arabidopsis (Bai et al., 1999).

Cohesin is key for chromosome segregation and its step-wise removal allows the segregation of homologous chromosomes in meiosis I, followed by separation of sister chromatids in meiosis II. In addition, cohesin is required for recombination and repair of DNA double-strand breaks resulting in a highly pleiotropic phenotype that leads to almost complete sterility of *rec8* mutant plants (Bai et al., 1999). Expression of our genomic *PRO_REC8_:REC8:GFP* reporter in a homozygous *rec8* mutant background completely restored fertility of these plants and analysis of chromosome spreads confirmed that chromosome segregation is indistinguishable from the wildtype (Supplement 1).

REC8 replaces RAD21 in meiosis and is hence highly specific for meiocytes in all species analyzed so far (Nasmyth, 2001). Consistent with previous immuno-localization studies, we found that the GFP signal of our functional reporter line was only present in meiocytes providing a straightforward way to identify microspore mother cells (Figure 2).

**Figure 2.**
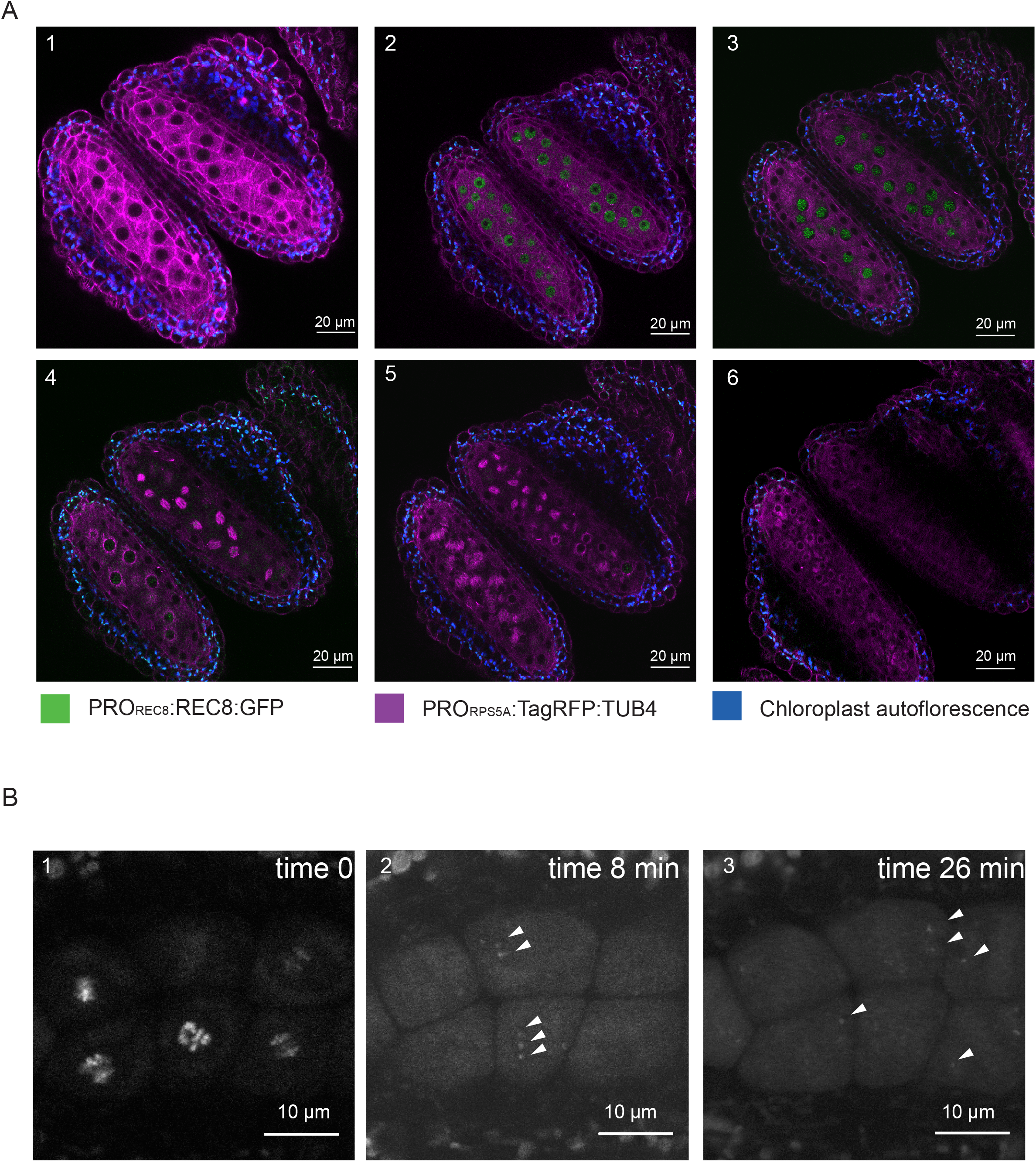
REC8 and TUB4 localization. A) A flower of the KINGBIRD line expressing PROREC8:REC8:GFP and PRORPS5A:TagRFP:TUB4 in meiosis. A1: premeiosis; A2: leptotene; A3: zygotene; A4: diplotene in the lower anther and metaphase I in the upper anther; A5: telophase I in the lower anther and late prophase Il-metaphase II transition in the upper anther; A6: tetrad stage. B) REC8-GFP localization after metaphase I (B1) in a PROREC8:REC8:GFP plant. The white arrowheads in B2 and B3 indicate centromeric REC8.

Moreover, the REC8 reporter allowed us to estimate the sensitivity of our imaging procedure. While REC8 is removed from chromosomes arms at the end of meiosis I to allow the resolution of cross-overs, a small fraction remains at the centromeres to maintain sister chromatid cohesion. The detection of the centromeric fraction of REC8 has been challenging by immuno-localization studies. When we followed the first meiotic division, we observed the remaining REC8 at centromeres indicating that our live cell imaging system is highly sensitive (Figure 2, Sup. movie 1).

Next, we combined the *PRO_rEC8_:REC8:GFP* with *PRO_RPS5A_:TagRFP:TUB4* that labels microtubules and hence permits observation of the cell shape and the formation of the spindles. The resulting double reporter line is referred to as *Kleisin IN Green microtuBules In ReD (KINGBIRD)* in the following. No meiotic defects and no reduction in fertility was observed in this line as well as in plants expressing only the *PRO_RPS5A_:TagRFP:TUB4* reporter (Supplement 1). The separated excitation and emission spectra of the two fluorochromes permitted the faithful and concomitant detection of both reporters.

An important question was how many frames per time interval should be taken. Due to photo-bleaching as well as potential photo-toxicity, a sampling rate of several frames per minute was not compatible with capturing the entire meiotic program. Based on a several test samples as well as previously published time courses (Armstrong et al., 2003; Sanchez-Moran et al., 2007; Stronghill et al., 2014; Yu et al., 1997), we decided to acquire one frame every 3 to 15 minutes, so that even the shortest phases such a metaphase I and II, can be captured while reducing photo-bleaching to a minimum.

**Sup movie 1: Detection of** *PROREC8:REC8:GFP* **at metaphase I –anaphase I transition**

This movie focuses on five male meiocytes at metaphase I. The *PROREC8:REC8:GFP* signal (in white) is seen on highly condensed chromosomes. With the onset of anaphase I, the remaining *PROREC8:REC8:GFP* can be seen at the centromere areas of homologs being pulled to opposite cell poles (white arrowheads at the movie reply).

**Supplement 1 Figure: Functionality of reporter lines used in this study.**

A) Stems with siliques of Col-0 WT, *rec8*, *REC8:GFP* in *rec8* background, and *KINGBIRD* in Col-0 WT background. While *rec8* has short and thin siliques (magenta arrowhead) indicating a high level of sterility, the *REC8:GFP* and *KINGBIRD* line are fertile as seen by their WT-like, elongated and thick siliques (white arrowheads).

B) Open siliques of Col-0 WT, *rec8*, *REC8:GFP* in *rec8* background, and *KINGBIRD* in Col-0 WT background. Col-0 WT, *REC8:GFP* and *KINGBIRD* form round, plumb and fully developed seeds (white arrow heads) whereas *rec8* has shriveled and small seed indicative of aborted seeds (magenta arrow head).

C) Anthers stained with Peterson staining. Col-0 WT, *REC8:GFP* and *KINGBIRD* only produce viable pollen while high rates of aborted pollen (blue) are visible inside the pollen sacs of *rec8* mutant plants.

D) Rate of seed abortion (%).

E) Rate of aborted pollen (%).

F) Chromosome spreads of Col-0 WT, *rec8, REC8:GFP* in *rec8* background, and *TagRFP:TUB4* in Col-0 WT background. The figure shows a selection of the main meiotic phases. Homozygous *rec8* mutants start to show defects from diplotene onwards, with the presence of univalent at metaphase I, and miss-segregating chromosomes afterwards leading to unbalanced tetrads and the formation of micronuclei. Wild-type meiotic progression is restored in *rec8* mutants expressing *REC8:GFP* and it is not disrupted in Col-WT plants expressing *TagRFP:TUB4*.

**Supplement 2: Table of primes used in this study**

### A meiotic landmark system

Meiosis is classically apportioned into nine phases: prophase I, metaphase I, anaphase I, telophase I, interkinesis, prophase II, metaphase II, anaphase II and telophase II. Due to the dramatic changes in chromatin structure and the dynamics of chromosomes, and to its prolonged duration, prophase I is divided into the five subphases leptotene, zygotene, pachytene, diplotene and diakinesis. These phases have been derived from observation of fixed material and chromosome spreads, leading to definitions mainly based on chromosome configuration, e.g. pachytene is defined by the presence of fully synapsed chromosomes.

Using the *KINGBIRD* reporter line, we were able to distinguish five parameters of meiocytes: cell shape, nucleus position, nucleolus position, microtubule array (MTs), and chromosome configurations (condensation and pairing/synapsis). Each parameter can adopt different states, e.g. the nucleus can be centrally located or in one corner of the meiocyte or absent (Figure 3A). Moreover, these parameter states have a distinct order, for instance we observed that cell shape always changed from rectangular to trapezoidal, to oval, to circular, to triangular and giving finally rise to tetrads composed of four triangular cells (Figure 3A). Importantly, our markers recapitulated previously described changes in nucleolus position and microtubule cytoskeleton, corroborating that our imaging system did not affect meiosis (Peirson et al., 1997; Stronghill et al., 2014; Wang et al., 2004).

**Figure 3.**
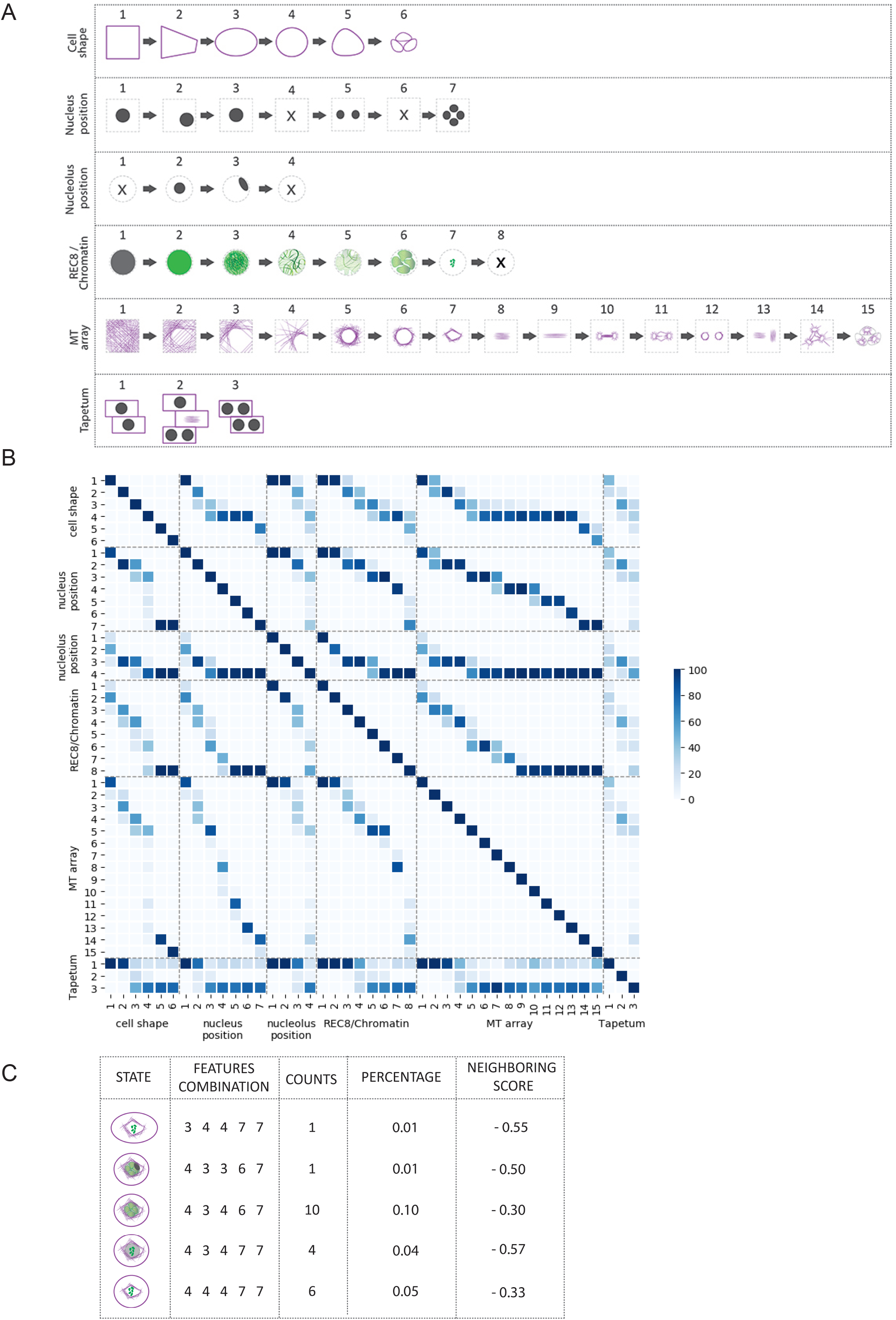
Identification of parameter states of male meiocytes. A) Schematic representation of the different states of the five parameters analyzed during meiotic progression. B) Heatmap of the correlation between the different states of the parameters. The darker the blue color, the tighter is the correlation and the higher is the frequency of co-appearance of two parameter states. Numbers refer to the scheme in A. C) Table illustrating the different parameter states at the moment of nuclear envelope breakdown. Even if the breakdown can be found (low number of observations), there is high variability of the combinations of parameter states that depict this moment. Hence, the neighboring scores are below zero, precluding the inclusion of the nuclear envelope breakdown as a landmark in this analysis.

Analyzing a first set of movies, gave rise to the hypothesis that some of the parameter states are connected, e.g. the nucleolus apparently dissolves only after the nucleus has moved to one side of the meiocyte and returned to a central position. To assess the nature of these associations, we analyzed a subset of cells (n=169 from 35 anthers) assigning a combination of numbers that represents each parameter state at every time point when a frame was taken, e.g. 1-1-2-2-1 describes a meiocyte that is rectangular in shape, has a centrally located nucleus with a centrally located nucleolus, with not condensed, yet not paired chromosomes and an evenly distributed microtubule array (Figure 3A). We subsequently analyzed 10,671 time points resulting from the first set of movies, that allowed us to judge which parameter states occur together and in which frequency (Figure 3B).

This analysis revealed that out of the more than 20,000 possible combinations of the different parameter states only 101 were actually present in our data set (Supplement 4) and their frequencies were distributed in a very dispersed range (from 0.01% to 21.14% of the total number of observations). In the following, we call a combination of all five parameter states a *cellular state*.

We realized at this point that an assessment of the cellular states, (e.g. concluding that the frequency of a state appearance in the dataset relates to its importance) is highly biased by the duration of the respective state, i.e. combinations of parameters that depict long phases such as pachytene are present in higher number of time points than combinations depicting short phases, e.g. metaphase I. Hence, to identify significantly distinct cellular states from the observed data, we defined a local or *neighboring score*, which quantifies the occurrence of a certain cellular state compared to its neighboring states.

A neighboring state was defined as a cellular state that is one transition away (−1 or +1) for at least one, but at most two, parameter states compared to the cellular state analyzed. With this, 2-2-3-4-4, for example, is a neighbor of 2-2-3-4-3 and of 3-2-3-4-3, but not of the cellular state 2-2-3-4-2 and not 3-2-3-3-3 (Supplement 4). Notably, we only took states into account that were actually observed. The neighboring score was then compared with the subset of *neighboring states*, to find the predominant state among the surrounding states, and is defined as:

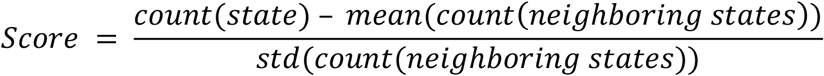

where *counts* refers to the number of times a certain state is observed in the data, and *std* refers to the standard deviation. This analysis revealed 11 clearly distinct cellular states that differed from their neighbors with a score higher than one, denoting that they occurred at least one standard deviation more frequent than the mean of the neighboring stages (Figure 4).

**Figure 4.**
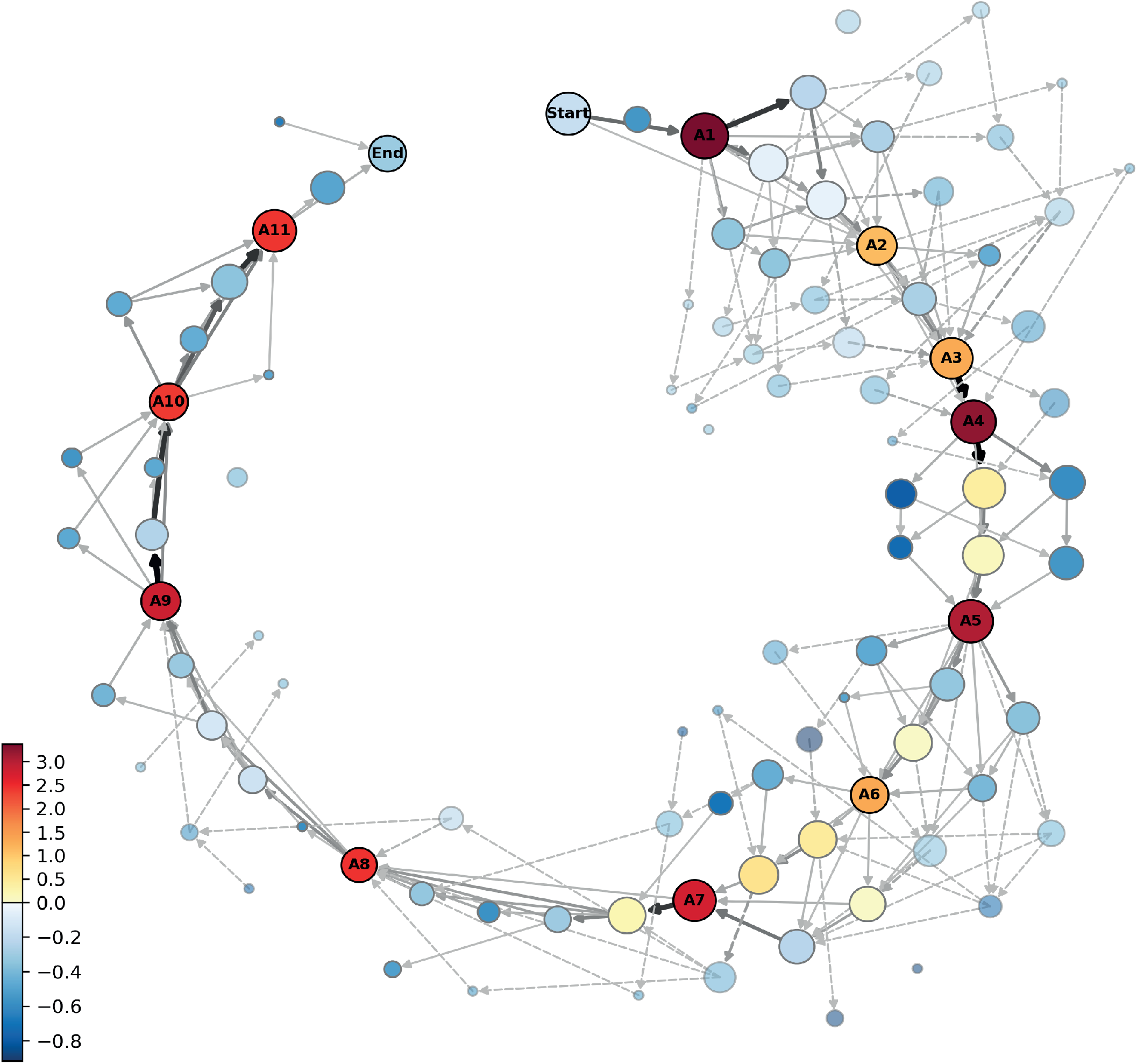
Meiotic landmarks. Meiosis represented as a progression of parameter state combinations, here called cellular states. Each circle signifies an observed cellular state and the arrows are observed transitions between these states. The size of circles depicts the frequency of appearance of each cellular state while the color presents their neighboring score. Cellular states that have a score higher than 1 are defined as landmarks and were assigned a name (A1-A11). Landmarks are highlighted by outlined circles and their names written in the center. The intensity of the line color of the arrows specifies which are the predominant paths taken by a male meiocyte undergoing meiosis. Notably the arrows indicate progression from one state to the following one only when the transition was seen within 15 minutes interval time, therefore the presence of non-connected circles.

These 11 cellular states (A1-A11) are henceforth called *meiotic landmark states* or *landmark* (Figure 4 and Figure 5). The states between landmarks are defined as *transition states*, and often represent alternative routes to the next landmark (Figure 4), e.g. the cell shape may first change from rectangular to trapezoidal and then the nucleus moves from a center position to a position at the side of the cell, or the nucleus moves first and then the cell shape changes. However, the nucleus is finally always located at the smaller side of the trapezoidal cell defining the new landmark state. The results of the neighboring score analysis were reproduced and confirmed by bootstrapping (Supplement 5).

**Figure 5.**
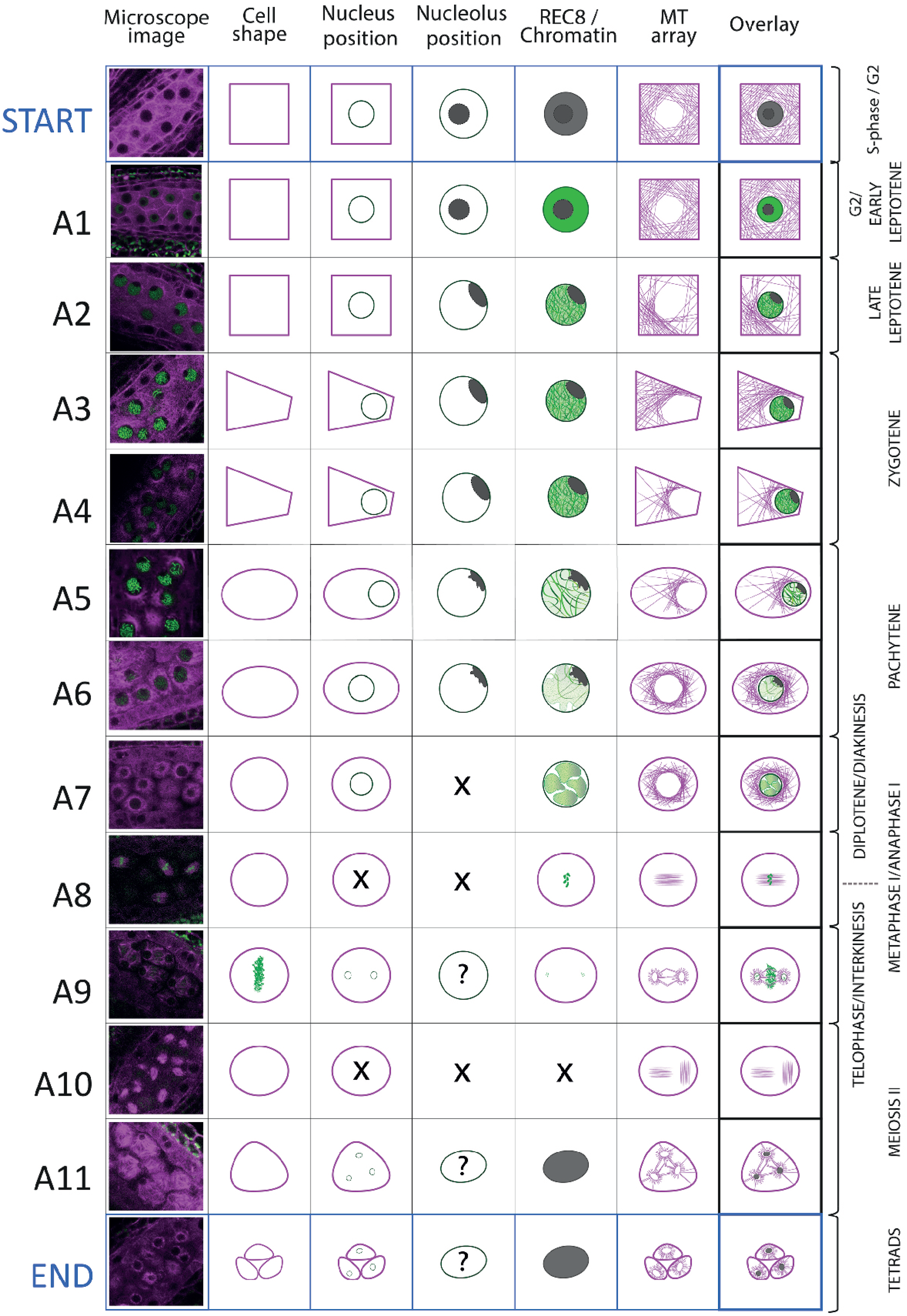
Landmark scheme. Illustration of the 11 here identified meiotic landmarks of male meiosis, A1-A11, and the combination of the parameter states that represent them. The first column provides a microscopy picture of meiocytes depicting each stage. The state of each parameter is separately shown in the following columns, the right-most column (Overlay) present their combination. On the right side, the classical stages of meiosis are assigned to each Landmark.

Taken together, we conclude that cellular differentiation steps of meiosis can be variable but then converge on distinct cell states, the landmarks. The qualitative assortment of the landmarks, possibly their order as well as their duration and the degree of variability (transition state number and duration), represent a new system to describe meiosis.

**Sup movie 2: Complete meiotic progression in the KINGBIRD line**

This movie follows two pollen sacs of different anthers from a single flower of the KINGBIRD line. Tubulin (RFP) is highlighted in magenta, chromosomes are marked by REC8 in green (GFP), chloroplasts (autofluorescence) are blue. The meiocytes, localized in the central areas of the pollen sacs, reside in a pre-meiotic stage at the beginning of the movie, and undergo a complete meiotic program with the first and the second meiotic division until the formation of tetrads. On the top left corner, there is an indication of the landmark crossed. Time in minutes; interval between image acquisition is 10 minutes, with the exception of 1 time point (7 minutes interval between time 1130 and 1137); time 0 corresponds to the start of image acquisition, and not to the start of meiosis.

**Supplement 3: Table of observed cell states**

**Supplement 4: Table summarizing the number of samples used per each analysis and the duration of each landmark**

**Supplement 5: Results of Bootstrap analysis**

### The case of the nuclear envelope breakdown

The break-down of the nuclear envelope in diplotene is an important hallmark of meiosis (Wijnker and Schnittger, 2013). We also could clearly observe the breakdown in our live cell imaging system although, due to its rapid progression, it was only captured in 22 out 10,671 analyzed time points with a sampling interval of one frame every 10 minutes (Figure 3C and Supplementary movie 3). None-the-less, the nuclear envelope break-down is not included in a landmark state since it appeared to be only loosely connected with the other parameter states, e.g. the cell shape can be oval or round, and the chromatin can be at different condensation levels when the nuclear envelope breaks down (Figure 3C). Thus, although very distinct when looking at microtubule conformation (i.e. state 7, collapse of pre-spindle Figure 3A), a clearly defined landmark state corresponding to nuclear envelope breakdown was not reached with the parameters analyzed.

We could also clearly observe other short-lived phases such as diakinesis, anaphase I, prophase II and anaphase II. However, due to their unexpected high variation in terms of association with the here analyzed parameter states, these phases, like nuclear envelope breakdown were also not designated as landmarks.

**Sup movie 3: NE breakdown in the KINGBIRD line**

This movie shows male meiocytes from diplotene to metaphase stage. The instance of nuclear envelope breakdown can be seen for the majority of the cells at minute 75 and for the remaining cells at minute 80. Tubulin is highlighted in magenta; chromosomes and REC8 are in green. The movie has been acquired with 5-minute interval time.

### Correlation between meiocyte and tapetum differentiation

Our sample preparation, which keeps anthers intact, also provided the possibility to follow the differentiation of the tissues surrounding the meiocytes, especially the tapetum cells. These are in direct contact with the meiocytes and are thought to nourish and support the meiocytes and spores (Pacini et al., 1985). A key feature of tapetum cells in many plants species, including Arabidopsis, is that they become poly-nucleated through endomitosis, i.e. a cell cycle variant in which cytokinesis is skipped (Jakoby and Schnittger, 2004). The poly-nuclearization of tapetum cells was clearly visible in our *KINGBIRD* line (Supplement movie 2, from minute 980 to minute 1207), possibly representing a sixth cell parameter next to the five meiotic parameters presented above (Figure 3A). Notably, tapetum cell differentiation was previously suggested as a criterion to judge stages of meiosis (Stronghill et al., 2014; Wang et al., 2004). We observed that polynucleated tapetum cells are not found before A4/zygotene and conversely, when all tapetum cells are poly-nucleated, meiosis has progressed into A7/diplotene. However, endomitosis only poorly correlated with any of the meiotic stages between A4 and A7 (Figure 3B) and hence, was not incorporated in the landmark system. In turn, we conclude that meiosis progresses largely independently of tapetum cell differentiation.

### Time course of meiosis in male meiocytes

The classical definition of meiosis, mostly relying on chromosome spreads, and the here-established landmark system by live cell imaging are based on different parameters and aspects of meiosis. For instance, chromosome spreads give a great spatial resolution of chromosome condensation levels that can currently not be reached in live cell imaging. Conversely, chromosome spreads loose several other cellular parameters such as the cytoskeleton and most crucially, do not allow following the behavior of a single meiocyte. None-the-less, the new landmark system described here can be assigned, albeit only roughly, to the classical phases of meiosis with A1 falling into an interval between S-phase and early leptotene, A2 in late leptotene, A3 and A4 in early and late zygotene, A5 and A6 in early and late pachytene, A7 in diplotene, A8 in metaphase I, A9 in interkinesis, A10 in metaphase II and finally A11 in telophase II (Figure 4). This led to the possibility to further test whether our imaging procedure might affect meiotic progression, determining the time course of meiosis in diploid Arabidopsis wild-type plants.

To estimate the duration of meiosis and approximate the landmarks to classical stages, the transition states between two landmarks were added to the time estimate of the preceding landmark. This also led to the re-assignment of diakinesis to A7, anaphase I to A8, prophase II to A9 and anaphase II to A10 (Figure 4). While long movies with more than 30 hours containing all meiotic stages could occasionally be obtained, they were rarely fully informative due to loss of the focal plane by sample growth. In contrast, 58 movies captured only subsections of meiosis, yet combined provided a complete coverage of meiosis I and II containing each landmark at least 4 times (Supplement 4). To faithfully judge the duration of each landmark, the length of one movie had to be long enough to capture at least two transitions of two sequential landmarks in one individual meiocyte (Sup. movie 2 and Supplement 4). We then tracked single meiocytes over time with up to 18 meiocytes per anther (Supplement 4).

Previously, the length of meiotic phases has been estimated by pulse-chase experiments in which either the modified thymine analog 5-bromo-2’-deoxyuridine (BrdU) or 5-ethynyl-2’-deoxyuridine (EdU) was applied to plants. After a given amount of time, meiotic spreads were prepared and tested for the appearance of these analogs in meiotic chromosome configurations (Figure 4). In these experiments, male meiosis in Arabidopsis was judged to last from G2 onwards approximately 32 to 33 hours with leptotene spanning between six and seven hours, zygotene and pachytene together lasting between 12 and 16 hours. Notably, these previous pulse-chase experiments were not able to resolve stages after diplotene and the rest of meiosis (from diplotene onwards) were estimated to approximately persist for three hours (Armstrong et al., 2003; Sanchez-Moran et al., 2007; Stronghill et al., 2014).

Our measurements of the meiotic phase lengths over all delivered a similar time frame and we determined the total duration of meiosis as 35 hours. This value includes the length of landmark A1 (8.5 hours in total), which stars with the onset of *REC8* expression, and therefore with S-G2 phase and ends at early leptotene stage. Prophase I, as expected, resulted to be the longest phase (minimum 20 hours) with late leptotene (A2) lasting 1.5 hours, zygotene (A3-A4) 6 hours, pachytene (A5-A6) 9.5 hours and diplotene and diakinesis (A7) together 3 hours. Importantly, we could also resolve meiotic phases thereafter and determine metaphase I and anaphase I (A8) together with 1 hour, telophase I, interkinesis and prophase II (A9) with 1 hour and meiosis II (A10-A11) all together with 4 hours (Figure 6).

**Figure 6.**
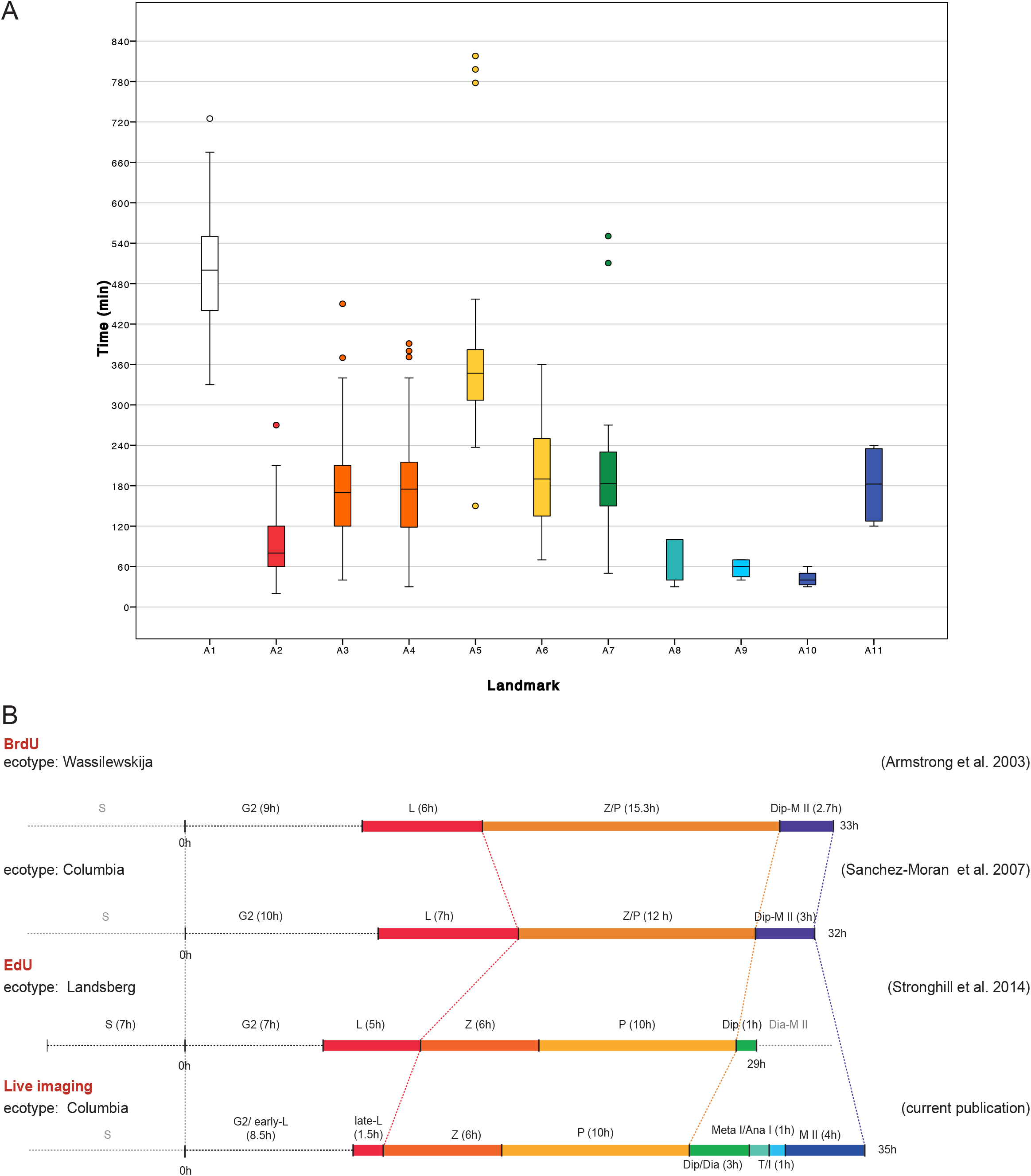
Time course of male meiosis in Arabidopsis. A) Box plot illustrating the duration of each landmark in minutes. Outliers are illustrated with a dot. The color code for each landmark refers to the meiotic phase: white (A1) is S-phase/G2, red (A2) is late leptotene, orange (A3, A4) is zygotene, yellow (A5, A6) is pachytene, green (A7) is diplotene/diakinesis, aquamarine (A8) is metaphase I/anaphase I, light blue (A9) is telophase /interkinesis, dark blue (A10, A11) is the second meiotic division. B) Comparison of meiotic timelines obtained with different techniques: BrdU and EdU staining, followed by sample fixation, versus live imaging. S stands for S-phase; L for Leptotene, Z for Zygotene, P for Pachytene, Dip for Diplotene, Dia for Diakinesis, Meta I/ Ana I for Metaphase and Anaphase I, T/I for Telophase and Interkinesis, M II for second meiotic division. The duration of each phase is indicated in hours for all the time courses, Time 0 has been set as the initiation of leptotene.

Summing up, the here-presented landmark system allows a dissection of meiosis with unprecedented temporal resolution. Given that the overall length of meiosis as well as the evaluation of individual subphases match previously determined durations, we conclude that our imaging system does not perturb meiosis and hence can be applied to analyze many mutant and to assess environmental conditions in the future.

**Supplement 4: Table summarizing the number of samples used per each analysis and the duration of each landmark**

## Discussion

Live cell imaging has promoted the understanding of many developmental and physiological processes. Examples from plants include live observation of double fertilization (Hamamura et al., 2014), following the division pattern in the shoot apical meristem (Gruel et al., 2016), the formation of leaf hairs (Bramsiepe et al., 2010), and tracking of cell death in the root (Fendrych et al., 2014). However, in contrast to following mitotic divisions and subcellular processes in the epidermis, live cell imaging of plant meiosis has been limited (Ingouff et al., 2017; Nannas et al., 2016; Sheehan and Pawlowski, 2009; Yu et al., 1997). The reasons for this are foremost the small number of meiocytes and their subepidermal position. In addition, appropriate reporters need to be introduced into the organism. Here, we have developed a robust *ex vivo* live cell imaging system for Arabidopsis male meiocytes based on conventional laser scanning microscopy. We show that this system does not lead to obvious alterations of meiotic progression when compared with previous time course analyses relying on pulse-chase experiments. Notably, our live imaging approach allows individual meiocytes to be followed; this overcomes the problem of asynchrony that occurs in late meiosis, which is likely the reason why the phases from diplotene to the end of meiosis II could not be resolved in the previous time course experiments. Thus, a more detailed judgment of short phases is a particular strength of our system resulting in a new powerful set up to analyze meiosis and describe the effects of mutants or environmental conditions.

### A landmark system

Our meiotic description is based on five morphological criteria of male meiocytes that we could distinguish with our reporter genes, i.e., cell shape, position of the nucleus, position of the nucleolus, REC8 status and information about chromatin state, and microtubule array. Importantly, we found that these cellular parameters have two aspects, which make them suitable for a classification system. First, they change in the course of meiosis in a unidirectional manner, e.g., cell shape changes from rectangular over trapezoidal and oval to circular. We never found an example where a meiocyte skipped one of these cell shape changes or changed back from a later stage to an earlier stage. Second, these parameters are linked with each other and build a matrix. For instance, nest-like microtubule array was never found to be associated with a rectangular cell shape of the meiocyte (Figure 3B).

Our analysis of cellular parameters allowed us to identify 11 prominent morphological states, called landmarks A1-A11. These differ from each other by at least one characteristic of the parameter states, and always occur in the same order in any cell progressing through meiosis. The pathway taken by an individual meiocyte to reach each landmark could differ slightly, presumably due to biological variation, and is described by the network of the transition states (Figure 4). It is an interesting question to what degree this developmental plasticity depends on meiotic genes and/or enviromental factors such as temperature.

The 11 landmarks together with their transitions could be assigned to the classical phases of meiosis (Figure 4). However, it has to be noted that the alignment of our landmarks with the classically defined stages remains fuzzy for certain phases. For example, leptotene is defined by the beginning of the chromosome pairing process, with the appearance of the first thin threads, a cell feature that we could not clearly resolve in our analysis. However, as more meiotic reporter lines are generated, for instance for the lateral or central elements of the synaptonemal complex, pairing and synapsis can be resolved with enhanced resolution in future. In this regard, the landmark system is highly modular and expandable depending on the resolution needed by the researcher.

Already with the current setup, our system allows an accurate and robust determination of meiotic stages. This is important since not all cell characteristics can always be unambiguously resolved, e.g. when the fluorescent signal diminishes because of photobleaching. Hence, the combined parameter states together with the knowledge about the previous cell stages maximize the information gained.

Our landmark system provides a powerful novel platform to study meiocyte differentiation and quantify meiotic progression. The observation that some of the cellular parameters are connected possibly indicates a common regulatory base and/or regulatory dependency. While some associations were expected, e.g. changes in MT cytoskeleton and cell shape, and are possibly directly linked, other combinations are new and unexpected, for instance the correlation between nucleolus movement and the MT cytoskeleton. These correlations can of course be indirect, yet exploring these combinations in future and identifying which genetic factors underly them opens a new perspective into meiosis. In turn, their potential uncoupling provides additional, qualitative criteria to describe meiotic mutants.

By adding more criteria in the future through the use of additional reporter lines and the analysis of mutants affecting meiosis, it will be possible to obtain a highly informative network of functional relationships, i.e. coupled and uncoupled parameters, within a meiocyte thus providing a system-biology understanding of meiosis. Importantly, it will be interesting to see to what degree these cellular parameters can be found and are coupled with each other in female meiocytes. Similarities and differences can further be compared with parameters of meiocytes in other plants species.

## Materials and Methods

### Plant material and growth conditions

The *Arabidopsis thaliana* (L.) Heynh plants used in this study were all derived from the Columbia (Col-0) ecotype. The *REC8 T-DNA* insertion line *rec8* (At5g05490, SAIL_807_B08) was obtained form Syngenta via NASC. All genotypes were determined by polymerase chain reaction (PCR) with the primers indicated in Table S1. All seeds were surface-sterilized with chloride gas, sown on 1% agar plates (half-strength Murashige and Skoog (MS) salts and 1% sucrose, pH 5.8) and stored 3 days at 4 °C in the dark for stratification. Antibiotics were added when required for seed selection: 25 mg/L Hygromycin B (Duchefa Biochemie B.V., Haarlem, The Netherlands). For germination, plates were transferred to long-day condition (16 h day/8 h night regime at 22 °C/18 °C). After germination, plants were transferred to soil and grown under short-day conditions for 2 weeks (12 h day/12 h night regime at 21 °C/18 °C), and then transferred to long-day conditions until seed production. For all crosses, flowers of the female parent were emasculated 1 day before anthesis and hand-pollinated 1 to 2 days later.

### Expression constructs: cloning and line selection

To generate the *PRO_reC8_:REC8:GFP* construct, a 7,145 bp genomic fragment of the *REC8* gene containing a 1.8 kbp fragment upstream of the ATG and 0.5 kbp fragment downstream of the stop codon was amplified with the primers AT5G05490-F and AT5G05490-R (TableS1) and cloned into pENTR/D-TOPO. A *SmaI* site was inserted in front of the stop codon of the *REC8* construct by PCR with the primers REC8 CterSmaI-F and REC8 CterSmaI-R (TableS1). The construct was linearized by *SmaI* digestion and ligated to a monomeric GFP (mGFP) fragment, followed by LR recombination reaction into the destination vector pGWB501 (Nakagawa et al., 2007)

A *REC8* reporter line was established by floral dip transformation of *rec8* heterozygous plants followed by selection of T1 plants on 0.5X MS agar plates supplemented with 25 mg/L Hygromycin B (Duchefa Biochemie B.V., Haarlem, The Netherlands) and 50 mg/L Carbenicillin. T2 seeds from individual T1 plants were germinated on 0.5X MS agar plates supplemented with 25 mg/L Hygromycin. T2 line #3 has been selected as the best performing line in terms of *rec8* rescued phenotype and fluorescence intensity.

The *PRO_RPS5A_:TagRFP:TUB4* line has been provided by Takashi Ishida, (Kumamoto University).

The *KINGBIRD* reporter line has been generated via crossing of plants containing the *PRO_REC8_:REC8:GFP* and *PRO_RPS5A_:TagRFP:TUB4* constructs as described above. Selection of transformant plants carrying both of the constructs has been done on Hygromycin plates for *PRO_REC8_:REC8:GFP* and screening for expression of *PRO_RPS5A_:TagRFP:TUB4* in root seedlings on plate (red filter, UV lamp, Olympus MVX10), before transferring on soil.

### Phenotype evaluation

Rescue of the *rec8* phenotype was assessed at pollen level using Peterson staining protocol as described in Peterson et al. (Peterson et al., 2010) and monitoring meiotic progression at a cytological level via cell spreads as described in Ross et al. (Ross et al., 1996).

### Live imaging of meiotic division

Flowers of 0.4-0.6 mm were isolated and prepared as presented in the results section *“Specimen preparation”*. Up to 4 samples were positioned on the same petri dish and cultured in Arabidopsis Apex Culture Medium (ACM): half-strength Murashige and Skoog (MS) salts, 1 % sucrose, 0.8 % agarose, pH5.8. Vitamins were added to a 1X concentration from 1000X stock solution (10 % Myoinositol, 0.1 % nicotin acid, 0.1 % pyridoxine hydrochloride, 0.1 % thiamine hydrochloride, 0.2 % gylicine dissolved in Millipore water and subsequentially filter sterilized) (Hamant et al., 2014). Time lapses were acquired using a Zeiss LSM 880 confocal microscope and ZEN 2.3 SP1 software (Carl Zeiss AG, Oberkochen, Germany). During image acquisition the petri dish was filled with autoclaved water and placed under a W-plan-Apochromat 40X/1.0 DIC objective (Carl Zeiss AG, Oberkochen, Germany). GFP was excited at λ 488 nm, and detected at λ between 498-550 nm. RFP was excited at λ 561 nm and detected at λ between 578-650 nm. Autofluorescence from chloroplasts was highlighted in blue using excitation at λ 488, and detection at λ between 680-750 nm. Time lapses were acquired as series of Z-stacks (6 planes, 50 μm distance). Interval time was varying from a max of 15 to a min of 3 minutes depending on sample conditions. The functions “Autofocus” and “Automatized positions” were used to acquire images. Room temperature and sample temperature were controlled and stabilized at 18 °C and 21°C respectively.

Image drift was corrected by the Stack Reg plugin (Rigid Body option) for Fiji. Cell numbers were assigned manually.

### Quantitative analysis of live cell imaging data

#### Data set description

The landmark system is based on the analysis of a subset of data on male meiocytes from WT plants carrying the KINGBIRD reporter constructs. A subset of the analyzable male meiocyte was described at every timepoint by assigning manually a value for each of the five parameters assessed. A total of 169 meiocytes from 35 anthers were annotated, leading to a total of 18,531 data points spanning more than 3,269 hours. For 7,860 observations one or more of the parameters could not be annotated with a well-defined state, with 5,893 observations not having a single parameter recognizable. The resulting dataset, consisting in 10,671 time points, was used to determine the co-occurrences of parameter states and the landmarks.

A second dataset was annotated solely using the landmark system for the comparison of the calculation of the time course. The timing of image acquisition and the landmark attribution for each cell at each time point was done manually. Landmarks were described using numbers from 0 to 12 with a trailing “s” when the landmark was appearing for the first time. An “n” was assigned to the cell when not visible at a certain time point. All the starting points appearing later than 15 min after the previous recorded landmark were discarded. Files in the CSV format were created for each anther.

All analysis of the two datasets were done using the Python programming language (Version 3.6, Python Software Foundation, https://www.python.org).

#### Landmark extraction: data preprocessing

The manually created data set contains a description of the state of each of the 5 parameters that were recorded in individual cells at 15 minute intervals. In some cases, the time between consecutive measurements was more than 15 minutes. In these cases, we inserted an unmeasured data point (‘n’) for each cell parameter such that time between measurements was equal and at most 15 minutes for each recorded time course. This was done to ensure that unmeasured periods are noted as unmeasured properly, and to decrease the risk to assign any unrealistic transitions. The combination of the state of each of the 5 cellular parameters makes up the cell state. Transitions from one cell state to another occur when one or more parameter transition to a new state.

#### Landmark extraction: Cell state co-occurrence

To create the co-occurrence heat map in Figure 4, we counted the number of times a combination of two parameter states occurred in the same cell at the same time point. Since some time courses were measured with different temporal resolution (e.g. 10 minute intervals versus 15 minute intervals), we first resampled the data points from all time courses to have the same time between measurements. Co-occurrence counts were normalized by the total number of counts in the column, including the counts where the state of the 2^nd^ parameter could not be measured. This means that in the upper triangular part of the matrix, the counts are normalized for the total occurrence of the first parameter (all parameter sections per row together sum up to 1) while in the lower triangular part of the matrix for the second parameter, each column sums up to 1.

#### Bootstrapping

To assess the robustness of the selected landmarks and thus our theoretical framework, we performed a bootstrapping procedure on our data set. The total set of observations was randomly sampled with replacement to obtain a data set 1.5 times the size of the original data set. Scores for each state in this data set were calculated using the procedure described in the previous paragraph. This process was repeated 1000 times to obtain estimates for the mean value, standard deviation and quantiles of the score of each cellular state.

### Meiotic time course calculation

The duration of each landmark was automatically extracted from the CSV files (Material and methods, *Quantitative analysis of live cell imaging data, Data set description*) using a custom software based on consecutive landmark transitions. This resulted in a dataset of 327 landmark durations from 136 meiocytes of 17 different anthers.

## Acknowledgments

We thank Chris Franklin (Birmingham University, UK) Maren Heese (University of Hamburg, DE) and Vanesa Calvo-Baltanas (Wageningen University and Research, NL) for critical reading and helpful comments to the manuscript. We are grateful to Olivier Hamant (ENS, Lyon) for training in imaging. The Versailles Arabidopsis Stock Center and the Nottingham Arabidopsis Stock Centre (NASC) are acknowledged for providing material used in this study. We thank Takashi Ishida (Kumamoto University, Japan) for providing us with the *PRO_RPS5A_:TagRFP:TUB4* construct.

## Author Contributions

M.A.P., S.K., E.W., and A.S. conceived and designed the experiments. M.A.P., S.K., K.M., and E.W., and performed the experiments. F.S. wrote a software for analysis of the data. A.S. contributed material and reagents. M.A.P., E.K., R.V.R., S.K., K.M., E.W., C.F. and A.S. analyzed the data. M.A.P., C. F. and A.S. wrote the article.

## References

Armstrong SJ, Franklin FCH, Jones GH. 2003. A meiotic time-course for Arabidopsis thaliana. Sex Plant Reprod 16:141–149. doi:10.1007/s00497-003-0186-4

Bai X, Peirson BN, Dong F, Xue C, Makaroff CA. 1999. Isolation and Characterization of SYN1, a RAD21-like Gene Essential for Meiosis in Arabidopsis. Plant Cell Online 11:417–430. doi:10.1105/tpc.11.3.417

Bramsiepe J, Wester K, Weinl C, Roodbarkelari F, Kasili R, Larkin JC, Hülskamp M, Schnittger A. 2010. Endoreplication controls cell fate maintenance., Endoreplication Controls Cell Fate Maintenance. PLoS Genet PLoS Genet 6, 6:e1000996–e1000996. doi: 10.1371/journal.pgen.1000996, 10.1371/journal.pgen.1000996

Crismani W, Girard C, Mercier R. 2013. Tinkering with meiosis. J Exp Bot 64:55–65.doi: 10.1093/jxb/ers314

Enguita-Marruedo A, Cappellen WAV, Hoogerbrugge JW, Carofiglio F, Wassenaar E, Slotman JA, Houtsmuller A, Baarends WM. 2018. Live cell analyses of synaptonemal complex dynamics and chromosome movements in cultured mouse testis tubules and embryonic ovaries. Chromosoma 1–19. doi: 10.1007/s00412-018-0668-7

Fendrych M, Van Hautegem T, Van Durme M, Olvera-Carrillo Y, Huysmans M, Karimi M, Lippens S, Guérin CJ, Krebs M, Schumacher K, Nowack MK. 2014. Programmed Cell Death Controlled by ANAC033/S0MBRER0 Determines Root Cap Organ Size in Arabidopsis. Curr Biol 24:931–940. doi:10.1016/j.cub.2014.03.025

Głuszek AA, Cullen CF, Li W, Battaglia RA, Radford SJ, Costa MF, McKim KS, Goshima G, Ohkura H. 2015. The microtubule catastrophe promoter Sentin delays stable kinetochore–microtubule attachment in oocytes. J Cell Biol 211:1113–1120. doi: 10.1083/jcb.201507006

Gruel J, Landrein B, Tarr P, Schuster C, Refahi Y, Sampathkumar A, Hamant O, Meyerowitz EM, Jönsson H. 2016. An epidermis-driven mechanism positions and scales stem cell niches in plants. Sci Adv 2:e1500989. doi: 10.1126/sciadv.1500989

Hamamura Y, Nishimaki M, Takeuchi H, Geitmann A, Kurihara D, Higashiyama T. 2014. Live imaging of calcium spikes during double fertilization in Arabidopsis. Nat Commun 5:4722. doi:10.1038/ncomms5722

Hamant O, Das P, Burian A. 2014. Time-Lapse Imaging of Developing Meristems Using Confocal Laser Scanning MicroscopePlant Cell Morphogenesis, Methods in Molecular Biology. Humana Press, Totowa, NJ. pp. 111–119. doi:10.1007/978-1-62703-643-6_9

Hand ML, Koltunow AMG. 2014. The Genetic Control of Apomixis: Asexual Seed Formation. Genetics 197:441–450. doi:10.1534/genetics.114.163105

Holubcová Z, Howard G, Schuh M. 2013. Vesicles modulate an actin network for asymmetric spindle positioning. Nat Cell Biol 15:937. doi:10.1038/ncb2802

Ingouff M, Selles B, Michaud C, Vu TM, Berger F, Schorn AJ, Autran D, Durme MV, Nowack MK, Martienssen RA, Grimanelli D. 2017. Live-cell analysis of DNA methylation during sexual reproduction in Arabidopsis reveals context and sex-specific dynamics controlled by noncanonical RdDM. Genes Dev 31:72–83. doi:10.1101/gad.289397.116

Jakoby M, Schnittger A. 2004. Cell cycle and differentiation. Curr Opin Plant Biol 7:661–669. doi:10.1016/j.pbi.2004.09.015

Kitajima TS, Ohsugi M, Ellenberg J. 2011. Complete Kinetochore Tracking Reveals Error-Prone Homologous Chromosome Biorientation in Mammalian Oocytes. Cell 146:568–581. doi:10.1016/j.cell.2011.07.031

Kyogoku H, Kitajima TS. 2017. Large Cytoplasm Is Linked to the Error-Prone Nature of Oocytes. Dev Cell 41:287–298.e4. doi: 10.1016/j.devcel.2017.04.009

Lambing C, Franklin FCH, Wang C-JR. 2017. Understanding and Manipulating Meiotic Recombination in Plants[OPEN]. Plant Physiol 173:1530–1542. doi:10.1104/pp.16.01530

Lambing C, Heckmann S. 2018. Tackling Plant Meiosis: From Model Research to Crop Improvement., Tackling Plant Meiosis: From Model Research to Crop Improvement. Front Plant Sci Front Plant Sci 9, 9:829–829. doi: 10.3389/fpls.2018.00829, 10.3389/fpls.2018.00829

Ma H. 2006. A Molecular Portrait of Arabidopsis Meiosis. Arab Book Am Soc Plant Biol 4. doi:10.1199/tab.0095

Mercier R, Mézard C, Jenczewski E, Macaisne N, Grelon M. 2015. The Molecular Biology of Meiosis in Plants. Annu Rev Plant Biol 66:297–327. doi: 10.1146/annurev-arplant-050213-035923

Mlynarczyk-Evans S, Villeneuve AM. 2017. Time-Course Analysis of Early Meiotic Prophase Events Informs Mechanisms of Homolog Pairing and Synapsis in Caenorhabditis elegans. Genetics genetics. 117.204172. doi: 10.1534/genetics. 117.204172

Mullen TJ, Wignall SM. 2017. Interplay between microtubule bundling and sorting factors ensures acentriolar spindle stability during C. elegans oocyte meiosis. PLOS Genet 13:e1006986. doi:10.1371/journal.pgen.1006986

Nakagawa T, Suzuki T, Murata S, Nakamura S, Hino T, Maeo K, Tabata R, Kawai T, Tanaka K, Niwa Y, Watanabe Y, Nakamura K, Kimura T, Ishiguro S. 2007. Improved Gateway Binary Vectors: High-Performance Vectors for Creation of Fusion Constructs in Transgenic Analysis of Plants. Biosci Biotechnol Biochem 71:2095–2100. doi:10.1271/bbb.70216

Nannas NJ, Higgins DM, Dawe RK. 2016. Anaphase asymmetry and dynamic repositioning of the division plane during maize meiosis. J Cell Sci 129:4014–4024. doi: 10.1242/jcs. 194860

Nasmyth K. 2001. Disseminating the Genome: Joining, Resolving, and Separating Sister Chromatids During Mitosis and Meiosis. Annu Rev Genet 35:673–745. doi: 10.1146/annurev.genet.35.102401.091334

Pacini E, Franchi GG, Hesse M. 1985. The tapetum: Its form, function, and possible phylogeny inEmbryophyta. Plant Syst Evol 149:155–185. doi: 10.1007/BF00983304

Peirson BN, Bowling SE, Makaroff CA. 1997. A defect in synapsis causes male sterility in a T-DNA-tagged Arabidopsis thaliana mutant. Plant J 11:659–669. doi:10.1046/j.1365-313X.1997.11040659.x

Peterson R, Slovin JP, Chen C. 2010. A simplified method for differential staining of aborted and non-aborted pollen grains. Int J Plant Biol 1:13. doi:10.4081/pb.2010.e13

Ross KJ, Fransz P, Jones GH. 1996. A light microscopic atlas of meiosis inArabidopsis thaliana. Chromosome Res 4:507–516. doi: 10.1007/BF02261778

Rosu S, Cohen-Fix O. 2017. Live-imaging analysis of germ cell proliferation in the C. elegans adult supports a stochastic model for stem cell proliferation. Dev Biol 423:93–100. doi:10.1016/j.ydbio.2017.02.008

Sanchez-Moran E, Santos J-L, Jones GH, Franklin FCH. 2007. ASY1 mediates AtDMC1-dependent interhomolog recombination during meiosis in Arabidopsis. Genes Dev 21:2220–2233. doi:10.1101/gad.439007

Schmidt A, Schmid MW, Grossniklaus U. 2015. Plant germline formation: common concepts and developmental flexibility in sexual and asexual reproduction. Development 142:229–241. doi:10.1242/dev.102103

Schuh M, Ellenberg J. 2007. Self-Organization of MTOCs Replaces Centrosome Function during Acentrosomal Spindle Assembly in Live Mouse Oocytes. Cell 130:484–498. doi: 10.1016/j.cell.2007.06.025

Sheehan MJ, Pawlowski WP. 2012. Chapter seven - Imaging Chromosome Dynamics in Meiosis in Plants In: P. Michael Conn, editor. Methods in Enzymology, Imaging and Spectroscopic Analysis of Living Cells Live Cell Imaging of Cellular Elements and Functions. Academic Press. pp. 125–143.

Sheehan MJ, Pawlowski WP. 2009. Live imaging of rapid chromosome movements in meiotic prophase I in maize. Proc Natl Acad Sci 106:20989–20994. doi: 10.1073/pnas.0906498106

Smyth DR, Bowman JL, Meyerowitz EM. 1990. Early flower development in Arabidopsis. Plant Cell Online 2:755–767. doi:10.1105/tpc.2.8.755

Stronghill PE, Azimi W, Hasenkampf CA. 2014. A novel method to follow meiotic progression in Arabidopsis using confocal microscopy and 5-ethynyl-2′-deoxyuridine labeling. Plant Methods 10:33. doi:10.1186/1746-4811-10-33

Wang Y, Wu H, Liang G, Yang M. 2004. Defects in nucleolar migration and synapsis in male prophase I in the ask1-1 mutant of Arabidopsis. Sex Plant Reprod 16:273–282. doi: 10.1007/s00497-004-0206-z

Wijnker E, Schnittger A. 2013. Control of the meiotic cell division program in plants. Plant Reprod 26:143–158. doi: 10.1007/s00497-013-0223-x

Yu H-G, Hiatt EN, Chan A, Sweeney M, Dawe RK. 1997. Neocentromere-mediated Chromosome Movement in Maize. J Cell Biol 139:831–840.

Zhou A, Pawlowski WP. 2014. Regulation of meiotic gene expression in plants. Front Plant Sci 5. doi:10.3389/fpls.2014.00413

